# Postnatal development of the dentate gyrus vascular niche

**DOI:** 10.1101/2025.04.21.649829

**Authors:** Nidhi Devasthali, India Carter, Angela I. Saulsbery, Elizabeth D. Kirby

**Affiliations:** Neuroscience Graduate Program, The Ohio State University, Columbus, OH, USA; Department of Psychology, College of Arts and Sciences, The Ohio State University, Columbus, OH, USA; Chronic Brain Injury Program, The Ohio State University, Columbus, OH, USA

**Keywords:** Dentate gyrus, hippocampal neurogenic niche, neural stem cells, blood vessels, postnatal development

## Abstract

Lifelong neurogenesis in the dentate gyrus (DG) of the hippocampus supports cognitive and emotional functions in most adult mammals. The subgranular zone (SGZ) of the DG contains dense vasculature where neural stem and progenitor cells (NSPCs) reside in close proximity to local capillaries. This arrangement likely supports NSPCs by providing access to oxygen, circulating molecules, and endothelial-derived factors. While SGZ vessel density and NSPC association with vessels are well established in adulthood, when these niche attributes emerge in development remains unclear. Here, we show that while blood vessel density in the SGZ remained stable from initial layer formation (2 weeks of age) into young adulthood (9 weeks of age) in male and female mice, the average distance from NSPC somas to the nearest blood vessel decreased progressively over postnatal development. This finding was accompanied by a symmetrical compression of proliferating cells within the SGZ, and a gradual shift of quiescent neural stem cell somas towards the granule cell layer of the DG. Our findings imply that the DG neurogenic vascular niche continues to refine postnatally, suggesting that the NSPC vascular niche has a unique functional role in supporting mature adult neurogenesis.

## Introduction

Adult neurogenesis occurs in two main neurogenic niches in the adult mammalian brain: the subventricular zone and the dentate gyrus (DG) of the hippocampus. In the mature DG, neurogenesis begins when radial glia-like neural stem cells (NSCs) in the subgranular zone (SGZ) divide to produce intermediate progenitor cells (IPCs), which subsequently mature into new neurons. This process of birth, survival, and maturation of new DG neurons is conserved across most mammalian species^1–3^. Studies in rodents have demonstrated that newborn DG neurons integrate in and modulate existing circuits, thereby supporting spatial learning and memory and affective behaviors^4–8^. Dysregulation of adult neurogenesis has been linked to neurological diseases such as Alzheimer’s disease and may even be a potential therapeutic target in other diseases and disorders with impaired hippocampal memory^6,9–11^.

In the adult DG, a major, essential feature of the stem cell niche is the local blood vessel network^12–15^. DG blood vessels originate as capillaries at the hippocampal fissure, expanding into a branched network in the molecular layer (ML). ML capillaries then converge and feed into capillaries that descend perpendicularly through the granule cell layer (GCL)^16^. These descending capillaries then turn 90° to run parallel to the SGZ, where they form a dense vascular network^16^. Within this dense, planar vasculature of the SGZ, NSCs (which are largely quiescent) and proliferating cells reside in close proximity to the blood vessels^14,17^. NSCs also wrap around blood vessels in the inner ML of the DG via their apical process^18,19^. This close association of neural stem and progenitor cells (NSPCs) to vessels may provide these neurogenic cells with heightened access to circulating oxygen, signaling molecules, and/or nutrient metabolites^16,20^. It may also uniquely expose neurogenic cells to endothelial-derived soluble factors which can inhibit differentiation, enhance neurogenesis, and promote self-renewal of NSCs^21,22^. Blood vessels may also serve as scaffolds guiding migration of IPCs and their differentiating progeny^17^.

Although the key features of vessel density and NSPC association with vessels are well characterized in the adult brain, the developmental progression of these features remains unclear. In rodents, the structure of the DG begins to develop prenatally from NSCs in the hippocampal primordium but is not complete until the second postnatal week when the layers of the DG are fully formed^23^. During the first postnatal week in mice, most of the granule neurons that will comprise the adult DG are generated from NSCs^23^. From early postnatal age, around postnatal day (PD) 17, to adulthood, around PD 60, there is a significant reduction in neurogenesis in the mouse DG^24^. This reduction in neurogenesis is accompanied by a decline in the number of proliferating cells during the same PD 17 to PD 60 time period^24^.

While the postnatal development of the neurogenic cell lineage has become better characterized in recent years, relatively little is known about how the local vasculature develops as these cell populations emerge and mature. Angiogenesis, the creation of new blood vessels from previously established ones, is generally considered complete in the brain by 3 weeks of age^25^. But these estimates are not specific to the unique DG neurogenic niche. When the vascular features of the DG layers arise and when the NSPC association with that vasculature is formed remain unclear. The DG vascular features and NSPC proximity to vasculature could develop in step with the formation of the DG and exhibit the adult phenotype as early as 2 weeks of age (when DG layers are first established), or conversely, they could continue to mature through postnatal development and into adulthood. To address this gap, we examined the vascular density across DG layers and the physical distance between NSPCs and local blood vessels from the juvenile age when DG layers are first formed (2 weeks) through young adulthood (9 weeks).

## Results

### Vascular density in the DG is dynamic over the course of postnatal development

To characterize changes in vascular density of each of the DG layers over postnatal development, we used immunolabeling for CD31 (a vascular endothelial marker) in brains harvested from male and female C57Bl/6J mice at 2, 3, 5 and 9 weeks of age (Fig. 1 A). These ages span from the juvenile period when the DG layers are first formed (2 weeks) to young adulthood (9 weeks). We measured percent area covered by CD31 immunoreactivity in the ML, GCL, SGZ, and hilus (HL) of the DG (Fig. 1 B, C). We found that vascular coverage did not change significantly between 2 and 9 weeks of age in the SGZ (all data reported as mean ± SEM; 2 weeks: 6.55 ± 0.78%; 3 weeks: 6.91 ± 0.33%; 5 weeks: 6.66 ± 0.37%; 9 weeks: 6.44 ± 0.36%), GCL (2 weeks: 2.30 ± 0.20%; 3 weeks: 2.45 ± 0.25%; 5 weeks: 2.53 ± 0.15%; 9 weeks: 2.44 ± 0.24%), and HL (2 weeks: 5.83 ± 0.70%; 3 weeks: 6.02 ± 0.45%; 5 weeks: 5.07 ± 0.33%; 9 weeks: 5.05 ± 0.36%). In the ML, vascular density decreased with postnatal age. CD31+ immunolabeled area at 9 weeks of age (5.80 ± 0.38%) was significantly higher than at 2 weeks (9.58 ± 0.74%), 3 weeks (9.37 ± 0.70%), and 5 weeks (7.89 ± 0.41%) of age. Put together, these findings suggest that the vascular density of most of the DG remains stable over postnatal development, with the exception of the ML which shows a moderate decrease in vascular density between juvenile and young adult ages.

**Fig 1:**
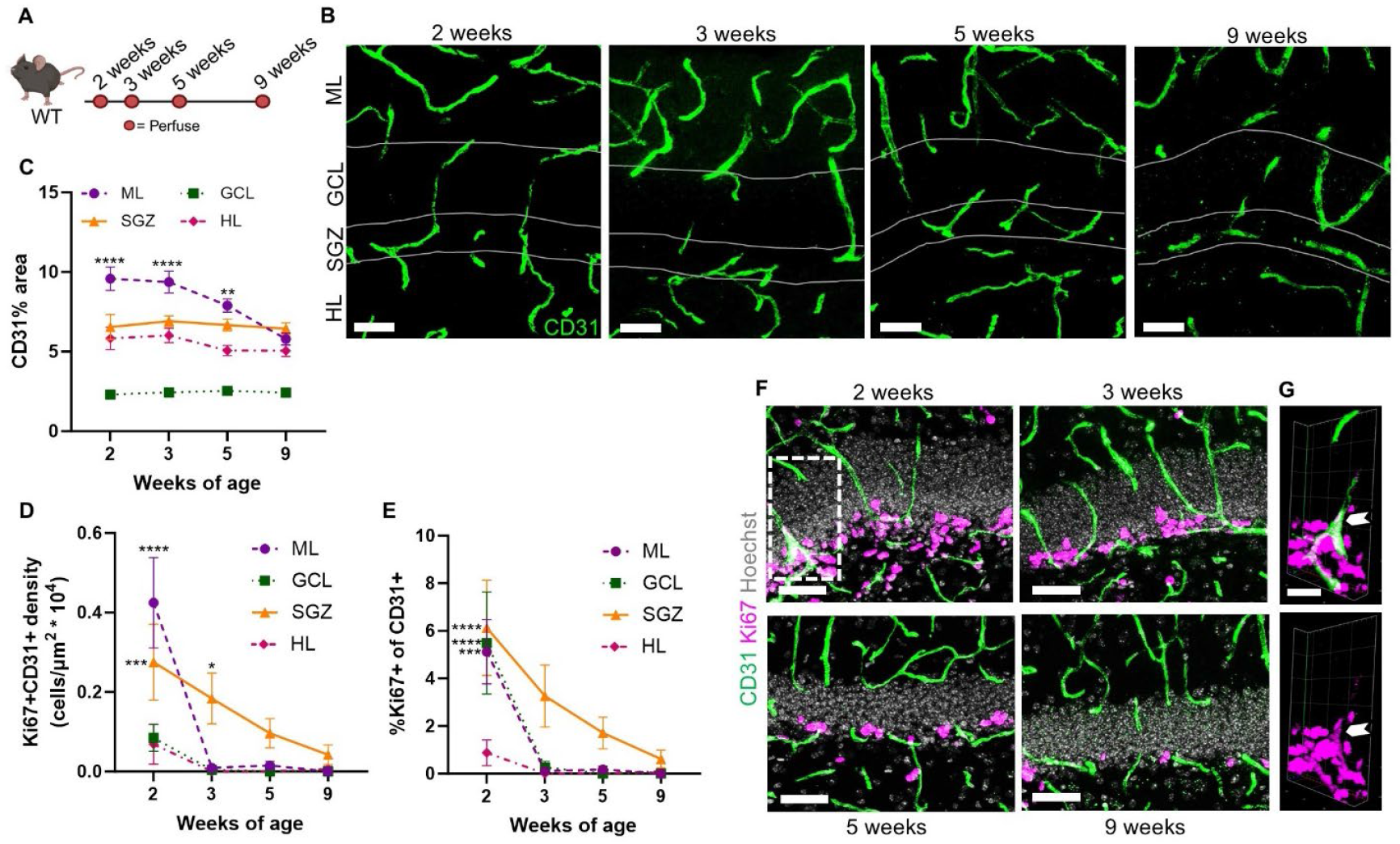
During postnatal development vascular coverage of the DG is stable in most layers and angiogenesis declines to near undetectable levels. **(A)** Experimental timeline to investigate the postnatal development of the DG neurogenic niche. **(B)** Representative images of immunolabeling for CD31. Scale = 50 µm. **(C)** Proportion of DG area covered by CD31 immunolabeling. 2-way repeated measures ANOVA age x layer (F_9,120_)=9.06, p<0.0001; age (F_3,40_)=2.20, p=0.10; layer (F_3,120_)=315.8, p<0.0001. **-p=0.0038, ****-p<0.0001 from Dunnett’s multiple comparisons to 9 weeks within layer. **(D)** Density of CD31+Ki67+ double labeled cells. 2-way repeated measures ANOVA age x layer (F_9,120_)=5.09, p<0.0001; age (F_3,40_)=11.28, p<0.0001; layer (F_3,120_)=12.41, p<0.0001. *-p=0.04; ***-p=0.0005; ****-p<0.0001 from Dunnett’s multiple comparisons to 9 weeks within layer. **(E)** Proportion of CD31+ cells co-labeled with Ki67. 2-way repeated measures ANOVA age x layer (F_9,120_)=2.27, p=0.02; age (F_3,40_)=12.60, p<0.0001; layer (F_3,120_)=8.39, p<0.0001. ***-p=0.0001, ****-p<0.0001 from Dunnett’s multiple comparisons to 9 weeks within layer. **(F)** Representative images of immunolabeling with CD31 and Ki67. Scale = 50 µm. **(G)** 3D representation of CD31+Ki67+ immunolabeling from area outlined in (F) with white arrowhead indicating CD31+Ki67+ double labeled cell. Scale = 20 µm. Mean ± SEM shown throughout with n = individual mice. 2 weeks (n=10), 3 weeks (n=12), 5 weeks (n=11) and 9 weeks (n=11).

### Angiogenesis in the DG is rare in postnatal development

We next combined CD31 immunolabeling with labeling for Ki67 (a cell cycle protein) to assess angiogenesis. While we were able to identify a small number of Ki67+CD31+ cells in all layers of the DG at 2 weeks of age (Fig. 1 D, F, G) (ML: 0.43 ± 0.11 cells/μm^2^ * 10^4^; GCL: 0.09 ± 0.03 cells/μm^2^ * 10^4^; SGZ: 0.28 ± 0.10 cells/μm^2^ * 10^4^; HL: 0.07 ± 0.05 cells/μm^2^ * 10^4^). By 3 weeks of age, Ki67+CD31+ cells were barely detectable in all DG layers (ML: 0.01 ± 0.01 cells/μm^2^ * 10^4^; GCL: 0.00 ± 0.00 cells/μm^2^ * 10^4^; SGZ: 0.18 ± 0.06 cells/μm^2^ * 10^4^; HL: 0.00 ± 0.00 cells/μm^2^ * 10^4^). Ki67+CD31+ cells were largely absent in all DG layers by 5 weeks of age (ML: 0.02 ± 0.01 cells/μm^2^ * 10^4^; GCL: 0.00 ± 0.00 cells/μm^2^ * 10^4^; SGZ: 0.10 ± 0.04 cells/μm^2^ * 10^4^; HL: 0.00 ± 0.00 cells/μm^2^ * 10^4^), a finding which was maintained at 9 weeks of age (ML: 0.00 ± 0.00 cells/μm^2^ * 10^4^; GCL: 0.00 ± 0.00 cells/μm^2^ * 10^4^; SGZ: 0.04 ± 0.03 cells/μm^2^ * 10^4^; HL: 0.01 ± 0.01 cells/μm^2^ * 10^4^). The rarity of proliferative CD31+ endothelia could also be appreciated when assessed as a proportion of CD31+ cells that were Ki67+ (Fig. 1 E). At their peak at 2 weeks of age, less than 7% of CD31+ cells were Ki67+ (ML: 5.12 ± 1.35%; GCL: 5.49 ± 2.15%; SGZ: 6.13 ± 2.00%; HL: 0.88 ± 0.54%). By 3 weeks of age, less than 4% of CD31+ cells were Ki67+ (ML: 0.10 ± 0.06%; GCL: 0.26 ± 0.26%; SGZ: 3.26 ± 1.30%; HL: 0.00 ± 0.00%), then less than 2% by 5 weeks of age (ML: 0.18 ± 0.13%; GCL: 0.00 ± 0.00%; SGZ: 1.71 ± 0.66%; HL: 0.00 ± 0.00%). By 9 weeks, the percent of CD31+ co-labeled for Ki67 was less than 1% (ML: 0.00 ± 0.00%; GCL: 0.00 ± 0.00%; SGZ: 0.61 ± 0.38%; HL: 0.10 ± 0.10%). Overall, these findings suggest that angiogenesis is rare in juvenile mice and declines to near or below detection limits in adulthood.

### Proliferative and quiescent NSPC soma proximity to vasculature increases over postnatal development

To examine proximity of NSPC somas to blood vessels over postnatal development, we quantified the shortest distance from proliferative and quiescent cell bodies in the SGZ to CD31+ blood vessel structures at 2, 3, 5, and 9 weeks of age (Fig. 2 A). We identified NSCs as SOX2+ cell bodies in the SGZ with a GFAP+ apical process extending through the GCL. Quiescent NSCs (qNSCs) were distinguished from proliferating NSCs by co-labeling for MCM2, a nuclear cell cycle protein (Fig. 2 B). The general proliferative population, which is primarily IPCs, was identified as Ki67+ nuclei in the SGZ. Quiescent and proliferative subsets of NSPCs all showed a similar pattern of being found closer to vessels as postnatal age advanced from juvenile ages (2 weeks) into young adulthood (9 weeks) (Fig. 2 C-G). Both MCM2- and MCM2+ NSCs were significantly closer to blood vessels at 9 weeks of age as compared to earlier time points (MCM2-NSCs: 2 weeks 17.35 ± 0.8526 µm; 3 weeks 14.08 ± 0.5590 µm; 5 weeks 12.20 ± 0.9253 µm; 9 weeks 11.78 ± 0.6851 µm; MCM2+ NSCs: 2 weeks 17.18 ± 0.8464 µm; 3 weeks 13.95 ± 0.8353 µm; 5 weeks 11.64 ± 1.327 µm; 9 weeks 10.27 ± 1.169 µm) (Fig. 2 E,F). Similarly, Ki67+ proliferative cells were significantly closer to vessels at 9 weeks of age than at 2 weeks of age (Ki67+ proliferative cells: 2 weeks 12.66 ± 0.5507 µm; 3 weeks 10.69 ± 0.2427 µm; 5 weeks 10.03 ± 0.4764 µm; 9 weeks 10.98 ± 0.4422 µm) (Fig. 2 G). Together, these findings suggest that NSC and IPC association with blood vessels continues to mature well into postnatal development.

**Fig 2:**
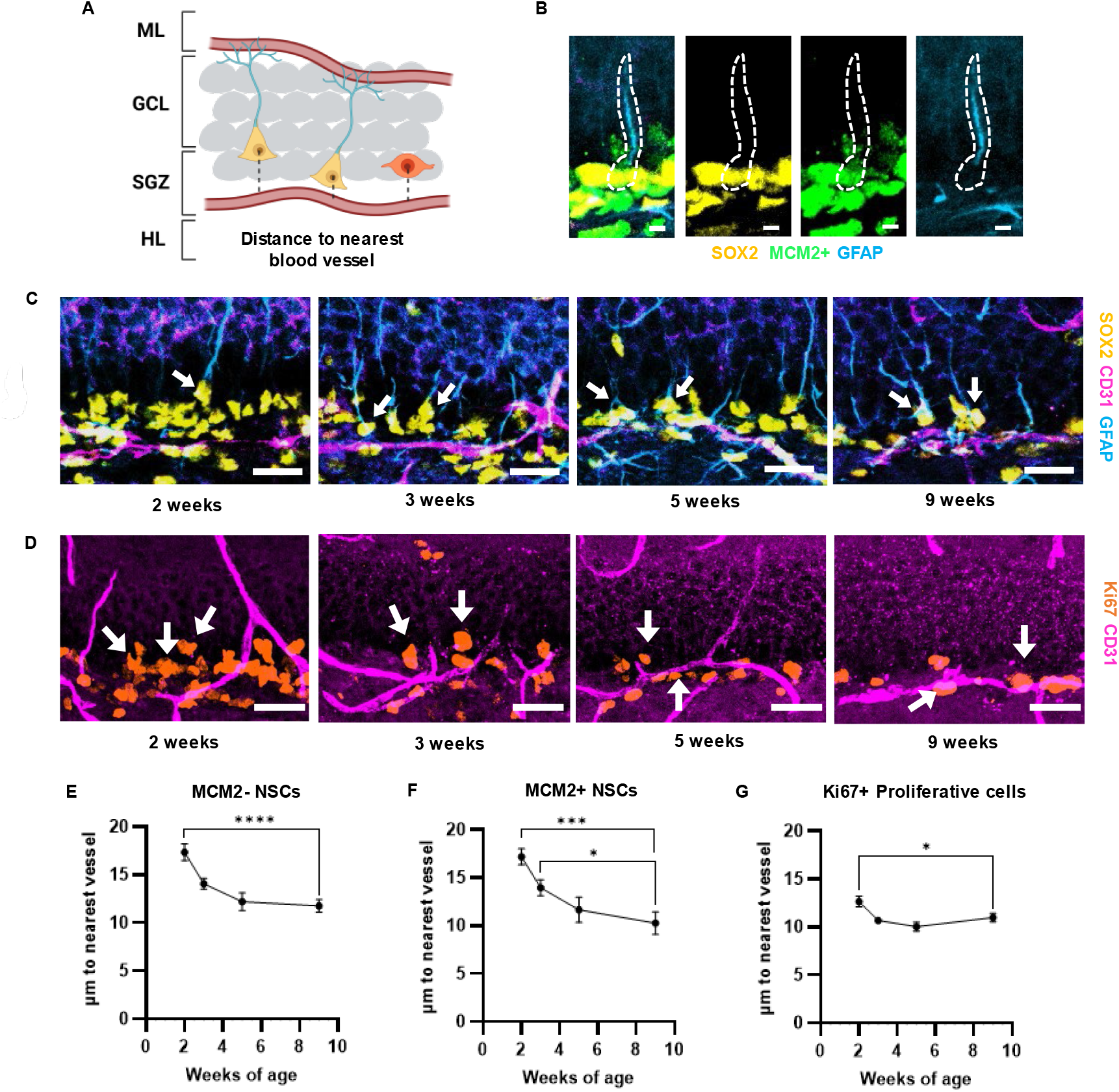
NSCs and proliferative cells in the DG get closer to blood vessels with age. **(A)** Graphic representation of the DG and the distance measurement taken from the center of the soma of NSCs and proliferative cells to the nearest blood vessel. NSCs are shown with yellow cell bodies and blue apical processes. Proliferative cells are shown as orange cell bodies. **(B)** Representative images of identification of an activated NSC using SOX2, MCM2, and GFAP immunolabeling. White outline is an identified NSC. Scale bar = 5 µm. **(C)** Representative images of CD31, SOX2, and GFAP immunolabeling. White arrows are pointing to identified NSCs. Scale bar = 20 µm. **(D)** Representative images of CD31 and Ki67 immunolabeling. White arrows are pointing to identified proliferative cells. Scale bar = 20 µm. **(E)** MCM2-NSC average distance to nearest CD31+ vessel. One-way ANOVA p<0.0001, F_(3,38)_ = 10.83, ****-p<0.0001 from Dunnett’s multiple comparisons to 9 weeks of age. **(F)** MCM2+ NSC average distance to nearest CD31+ vessel. One-way ANOVA p=0.0003, F_(3,38)_ = 7.936. ***-p=0.0002, *-p=0.0413 from Dunnett’s multiple comparisons to 9 weeks of age. **(G)** Ki67+ proliferative cell average distance to nearest CD31+ vessel. One-way ANOVA p=0.0011, F_(3,38)_ = 6.556. * - p=0.0264 from Dunnett’s multiple comparisons to 9 weeks of age. Mean ± SEM shown throughout with n = individual mice. 2 weeks (n=10), 3 weeks (n=12), 5 weeks (n=10) and 9 weeks (n=10).

### Quiescent NSCs become located closer to the GCL over postnatal development

To further investigate the NSPC niche throughout postnatal development, we quantified the position of quiescent and proliferative NSPCs within the DG relative to the midline of the SGZ at 2, 3, 5 and 9 weeks of age by measuring the shortest distance from the center of NSPC somas to the midline of the SGZ. Distances toward the GCL were assigned positive values and distances toward the HL were assigned negative values (Fig. 3 A). We found that the mean location of GFAP+SOX2+MCM2- (quiescent) NSC somas shifted significantly toward the GCL over postnatal development (Fig 3 A, B). The MCM2-NSCs were significantly closer to the GCL at 9 weeks of age as compared to both 2 and 3 weeks of age (MCM2-NSCs: 2 weeks -0.1503 ± 0.6201 µm; 3 weeks 0.6368 ± 0.4042 µm; 5 weeks 2.436 ± 0.5384 µm; 9 weeks 3.084 ± 0.5020 µm) (Fig. 3 B). The general pattern seen in the mean soma location of the MCM2-NSCs was not found in either proliferative cell population (MCM2+ NSCs: 2 weeks 0.2577 ± 0.9202 µm; 3 weeks -0.3914 ± 0.6713 µm; 5 weeks 0.3959 ± 0.6227 µm; 9 weeks 1.548 ± 0.9527 µm, Ki67+ proliferative cells: 2 weeks -0.7986 ± 0.5421 µm; 3 weeks -2.276 ± 0.4840 µm; 5 weeks -1.784 ± 0.2807 µm; 9 weeks -0.1159 ± 0.5567 µm) (Fig. 3 C, D). To look more closely at each cell type’s spatial distribution in the DG, we used violin plots to visualize cell soma location distribution relative to the SGZ midline. As expected based on the mean cell soma location data, the majority of cells in the MCM2-NSC population were found gradually closer to the GCL border of the SGZ with age (Fig. 3 E). The MCM2+ NSC population, in contrast, was spread more evenly across the SGZ and this distribution was relatively stable across ages (Fig. 3 F). The Ki67+ proliferative population was also centered in the SGZ across ages, though it became more concentrated around the SGZ midline throughout postnatal development as the population of Ki67+ qualitatively dwindled (Fig. 3 G). All together, these results suggest that the physical location of NSPCs in the local DG niche continue to mature throughout development.

**Fig 3:**
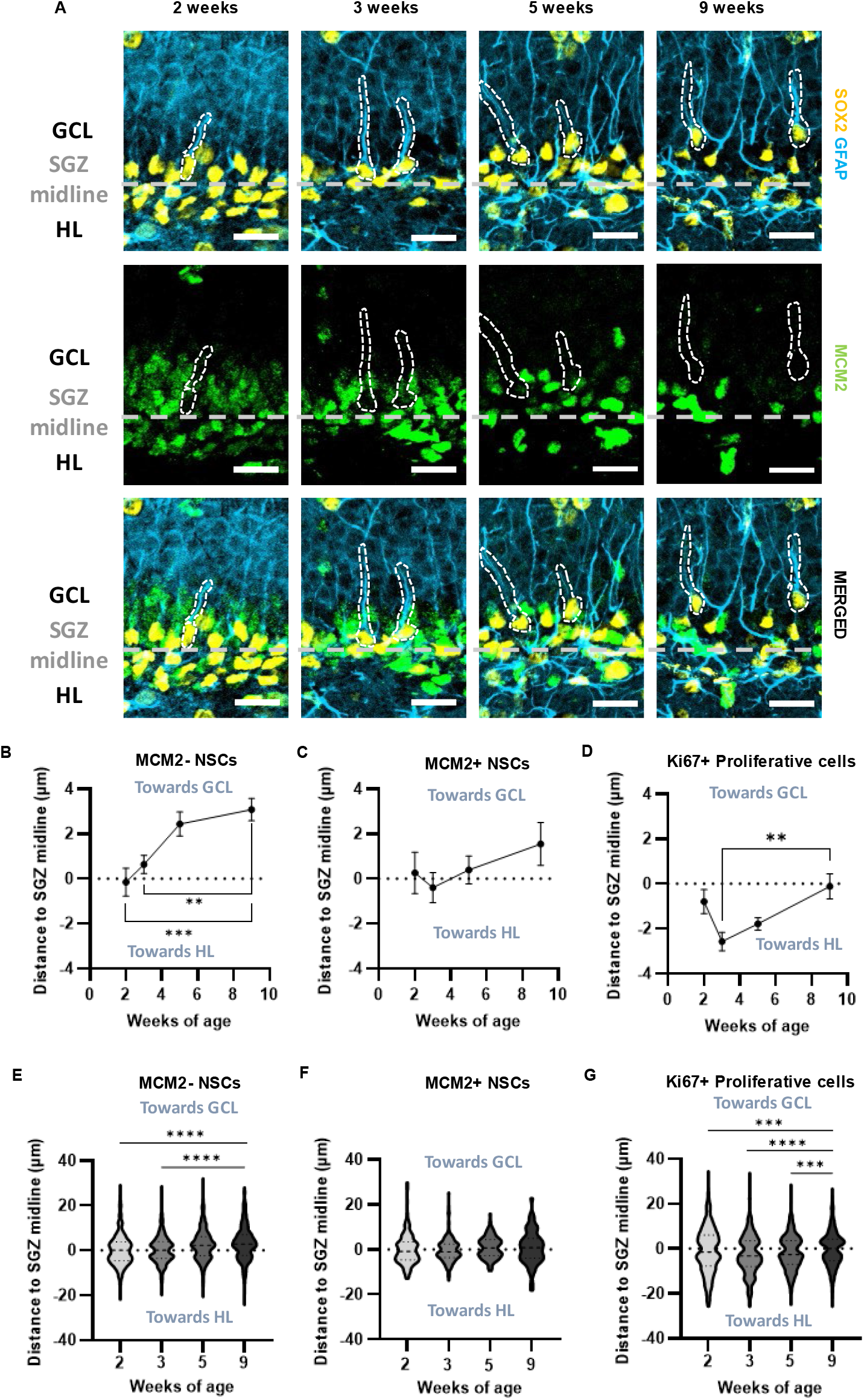
Quiescent NSCs in the DG are located closer to the GCL with age. **(A)** Representative images of SOX2, GFAP, and MCM2 immunolabeling. The SGZ midline is denoted by a dashed grey line with the GCL above and HL below. White outlines are identifying quiescent MCM2-NSCs. Scale bar = 20 µm. **(B)** MCM2-NSC average distance to SGZ midline. One-way ANOVA p=0.0002, F_(3,38)_ = 8.447. ***-p=0.0003, **-p=0.0040 from Dunnett’s multiple comparisons to 9 weeks of age. **(C)** MCM2+ NSCs average distance to SGZ midline. One-way ANOVA p=0.3820, F_(3,38)_ = 1.049. **(D)** Ki67+ proliferative cells average distance to SGZ midline. One-way ANOVA p=0.0121, F_(3,38)_ =4.158. **-p=0.0069 from Dunnett’s multiple comparisons to 9 weeks of age. **(B – D)** Mean ± SEM shown throughout with n = individual mice. 2 weeks (n=10), 3 weeks (n=12), 5 weeks (n=10) and 9 weeks (n=10). **(E)** Violin plots of distance to SGZ midline from individual MCM2-NSCs. p<0.0001 between 9 and 2 weeks and p<0.0001 between 9 and 3 weeks of age from Kolmogorov-Smirnov test. 2 weeks (n= 632 cells), 3 weeks (n=838 cells), 5 weeks (n=763 cells), and 9 weeks (n=760 cells). **(F)** Violin plots of distance to SGZ midline from individual MCM2+ NSCs. 2 weeks (n=181 cells), 3 weeks (n=192 cells), 5 weeks (n=85 cells), 9 weeks (n=55 cells). **(G)** Violin plots of distance to SGZ midline from individual Ki67+ proliferative cells. p=0.0002 between 9 and 2 weeks, p<0.0001 between 9 and 3 weeks, and p=0.0001 between 9 and 5 weeks of age from Kolmogorov-Smirnov test. 2 weeks (n=856 cells), 3 weeks (n=1043 cells), 5 weeks (n=892 cells), 9 weeks (n=847 cells).

## Discussion

The hallmark features of the DG neurogenic vascular niche are widely hypothesized to be critical for supporting lifelong neurogenesis. Here, we show that while some aspects of the vascular structure of the DG are present when the DG layers are first formed at 2 weeks of age in mice, the vascular niche continues to mature throughout postnatal development into young adulthood. Specifically, the relative vascular density of the DG ML decreases while other layers remain stable, and NSPC soma-vessel association progressively increases through postnatal development.

During embryonic development, the majority of brain vascularization is achieved via angiogenesis, the division of existing vascular endothelial cells to create new blood vessels^26^. Angiogenesis is generally considered complete by 3 weeks of age in the mouse brain, though this data is mostly based on measurement of cortical tissue^27^. Consistent with these previous findings, we found almost no evidence of angiogenesis in the DG beyond 2 weeks of age, as reflected by the scarcity of CD31+ endothelial cells expressing the cell cycle marker Ki67. We further found that the vascular density in the GCL, SGZ, and HL layers of the DG niche remained constant from juvenile ages to young adulthood. The maintenance of vascular coverage along with the absence of angiogenesis suggests that vascular endothelial cells in the GCL, SGZ and HL likely elongate or widen the diameter of blood vessels to ensure consistent vascular coverage as total brain volume increases during postnatal development. The ML layer of the DG, in contrast, lost vascular density between 2 and 9 weeks of age. Why the ML loses vascular coverage while other DG niche layers do not, remains unclear.

In the adult mammalian DG, the close proximity of NSPC bodies to blood vessels in the SGZ has 2 major hypothesized functions: 1) to bring NSC progeny in proximity to vessels that they will use as scaffolding for migrating along as they differentiate^17^, and 2) to provide specialized access to blood and/or endothelial signals that support their survival and proliferation^16,20,22,28^. Our data show the average distance between blood vessels and NSPC somas decreased over juvenile development into young adulthood, suggesting an increasing association of NSPCs to vessels as the DG matured. Proliferative cell somas maintained relatively stable locations within the SGZ, though the width of the proliferative zone compressed with age. The progressive increase in average association of proliferative NSPCs with local vessels suggests functional importance of this relationship. Quiescent NSCs also showed increasing proximity to SGZ vessels with age, coupled with a shift in the location of their somas within the SGZ to be on average closer to the vascularly sparse GCL at older ages than at younger ages. This combination of concentrating in a generally less vascular area while increasing average vessel association strongly suggests that quiescent NSC proximity to blood vessels carries functional relevance.

Our data do not establish the cellular or molecular mechanisms by which NSPC-vessel proximity develops postnatally. One possible cellular mechanism explaining how NSPC somas become closer to vessels on average over time is through the selective maintenance or proliferation of NSPCs closest to blood vessels. Over the course of postnatal development, the number of NSPCs decreases dramatically^29,30^. This decrease is particularly notable in the proliferative cell population as NSCs become progressively more quiescent and IPCs and their progeny differentiate^31,32^. If proximity to blood vessels promotes maintenance and/or proliferation of NSPCs, then the loss of NSPCs farther away from blood vessels would decrease the average NSPC-vessel distance over time. An additional possible mechanism for increased NSPC soma association with vessels over postnatal development is that NSPCs may migrate towards blood vessels as the niche matures. Finally, a third possibility is that vessels migrate towards NSPCs. All three of these cellular mechanisms could be operating simultaneously or in any combination to drive greater NSPC-vessel association. Ultimately, further investigation into the mechanisms driving the development of NSPC-vessel proximity is needed to determine how the association develops.

The molecular signals that direct increased postnatal NSPC-vessel association are also unclear. Broadly speaking, blood-derived signals seem to be likely candidates which could promote NSPC survival and proliferation near vessels, and/or migration towards vessels. For example, growth factors like insulin-like growth factor and brain-derived neurotrophic factor are found in the blood and promote NSPC proliferation and survival embryonically and during adulthood^33–37^. Blood-borne signals have been shown to drive adult neurogenesis changes in response to systemic stimuli like exercise^36,38–41^ and aging^22,28,42,43^, and this type of blood-borne regulation may extend to this postnatal refinement of the neurogenic vascular nice. NSPC-derived signals also could play a role in postnatal maturation of NSPC-vessel association. For example, NSPCs secrete a variety of factors that could affect their own migration or the physiology of local blood vessels^44–47^. Most notably, NSPCs synthesize vascular endothelial growth factor (VEGF)^48,49^. VEGF is required for normal vascularization of the embryonic brain, with endothelial cells growing towards the VEGF secreted by embryonic NSCs^50,51^. In adulthood, we recently showed that DG NSPC-derived VEGF is essential for maintaining NSPC-vessel association^52,53^. Surprisingly, this dependence appeared to rely on self-stimulated NSPC motility and cell attachment, rather than changes in vascularization. In the current study, NSPC-VEGF could play a role in vessels migrating towards NSPCs and/or maintaining existing NSPC-vessel proximity. More work is needed to define the molecular signals that guide postnatal vascular niche maturation.

There are several limitations to our study. First, while we restricted our investigation of NSPC-vessel association to the SGZ, previous literature has demonstrated that NSC processes wrap around blood vessels in the ML, and potentially gain access to blood factors at these contact points^18,19^. Exploring NSC-vessel wrapping at functional and structural levels during postnatal development, a time when the vascular density of the ML in particular is changing, could lend greater insight into the hypothesized functional importance of vessel wrapping on NSC survival and proliferation. Second, our analysis of vascular coverage by DG layer reflects broad changes in vascular patterns, but would be unable to reveal any subtle movements of vessels within DG layers. Studies using live imaging of blood vessels could identify vasculature migration patterns, and determine if migration of blood vessels contributes to increasing NSPC-vessel association. Third, our analysis of NSPC-vessel proximity used 2-dimensional images to determine the distance between the center of NSPC cell somas and the nearest blood vessel. These methods are in keeping with standards of the field^52^, in particular with the original study establishing the close association of proliferative cells with blood vessels^14^. These methods provide a useful approximation of NSPC-vessel association, but they lack 3-dimensional detail by definition. Further studies using 3-dimensional imaging could lend new insight into the intricacies of NSPC-vessel interactions.

In conclusion, we have found that the DG neurogenic niche continues to mature through postnatal development. Our characterization of the DG niche during juvenile development forms a framework for future investigation into the mechanisms guiding the changes we observed. Identifying the mechanisms of niche development and determining their functional relevance would ultimately help better understand how the hallmark features of the adult DG neurogenic niche arise. Understanding the development of key niche features could also aide in creating microenvironments receptive to stem cell transplants, paving the way for treatments for several neurological disorders and diseases.

## Methods

Authors complied with ARRIVE guidelines while conducting and reporting experiments described here.

### Mice

Timed pregnant wild-type C57BL/6J female mice (n = 7) were obtained from Jackson labs (#000664) and were housed individually with a toy in The Ohio State University Psychology Building mouse vivarium. All mice were housed in standard ventilated cages on a 12/12 hour light/dark cycle with *ad libitum* access to food and water. The pups were weaned at 3 weeks of age and housed in same sex groups of 2-5. All animal use was in accordance with institutional guidelines approved by The Ohio State University Institutional Animal Care and Use Committee.

### Experimental Design

Mice were randomly selected from 7 litters at 2 weeks (n = 10, 5M/5F), 3 weeks (n = 12, 8M/4F), 5 weeks (n = 11, 5M/6F), and 9 weeks (n = 11, 5M/6F) of age. Each timepoint had mice from at least 6 of the 7 litters. No mice with viable immunolabeling were excluded from final analyses. Immunolabeling was insufficiently clear or absent in one or more channels for 2 mice in the GFAP/SOX2/MCM2/CD31 immunolabeling only (1 mouse from 5 weeks of age group, 1 mouse from 9 weeks of age group). All mice received intraperitoneal injections of ethynyl-deoxyuridine (Click Chemistry Tools, #1149-500), dissolved fresh in sterile physiological saline at 10 mg/ml at dosage 150 mg/kg 2 hours before being perfused. At the indicated tissue harvest time, mice were anesthetized with a mixture of 87.5 mg/kg ketamine and 12.5 mg/kg xylazine, then transcardially perfused with 0.1M phosphate buffered saline (PBS).

### Tissue Processing

Mouse brains were post-fixed with 4% paraformaldehyde in 0.1M PBS at 4°C for 24 hours. They were then equilibrated in 30% sucrose in 0.1 M PBS at 4°C. Brains were then sliced on a freezing microtome in a 1:12 series of coronal sections at 40 µm thick. Sections were stored in cryoprotectant medium at -20°C until used for immunofluorescent staining.

### Immunolabeling

For Ki67 and CD31 immunolabeling, free-floating sections were rinsed in PBS, blocked in 0.3% triton-100X and 1% normal donkey serum in PBS for 30 min at room temperature, then incubated in primary antibody in blocking solution overnight on a shaker at 4°C (Table 1). The next day, the sections were rinsed with PBS then incubated in secondary antibodies (Table 1) in blocking solution for 2 hours on a shaker at room temperature. The sections were then labeled with Hoechst 33342 (1:2000 in PBS, Fisher #H3570), followed by final rinses in PBS.

**Table 1.**

| Table 1 |  |  |  |  |  |
| --- | --- | --- | --- | --- | --- |
| Primary antibody | Vendor/product no | Dilution | Secondary antibody | Vendor/product no | Dilution |
| Rat anti-CD31 | BD Pharmingen 550274 | 1:100 | Donkey Alexafluor594 anti-rat | Fisher A21209 | 1:500 |
| Rabbit anti-Ki67 | Cell Signaling Technology 9129 | 1:400 | Donkey Alexafluor647 anti-rabbit | Fisher A31573 | 1:500 |
| Rat anti-SOX2 | Affymetrix eBioscience 14-9811 | 1:1000 | Fab fragment Goat Alexa Fluor594 | Jackson Immuno 112-587-008 | 1:500 |
| Mouse anti-GFAP | EMD Millipore MAB360 | 1:1000 | Donkey Alexa fluor350 antimouse | Fisher A10035 | 1:1000 |
| Rabbit anti-MCM2 | Cell Signaling Technology 4007 | 1:500 | Donkey Alexa fluor647 antirabbit | Fisher A-31573 | 1:500 |

For GFAP/SOX2/CD31/MCM2 labeling, slices were first labeled for SOX2 as above, followed by a 10 min fixation step in 4% paraformaldehyde at room temperature. Then they were labeled for GFAP/CD31/MCM2 following the protocol above.

Immunolabeled sections were mounted on SuperFrost Plus slides, dried overnight at room temperature, and cover slipped with Prolong Gold Antifade mounting medium. For long term storage, slides were placed in the dark at 4°C.

### Microscopy

The DG was imaged at 20x magnification in 15 µm z-stacks with 1 µm steps using Zeiss Axio Observer Z.1 with apotome digital imaging system and axiocam 506 monochrome camera (Zeiss).

### Image Quantification

Layers of the DG for all image quantification were defined as – ML: the region between the suprapyramidal blade of the granule cell layer and the stratum lacunosum-moleculare of the CA1 region combined with the region between the infrapyramidal blade of the granule cell layer and the end of the hippocampal structure; GCL: the dense layer made up of cell bodies of granule cells excluding the SGZ; SGZ: the zone spanning 2 cell body widths between the granular cell layer and the hilus; HL: the region between the suprapyramidal and infrapyramidal blades of the granule cell layer, excluding the SGZ and CA3 regions.

For vascular coverage analysis, layers of the DG (ML, GCL, SGZ, and HL) were outlined in maximum intensity projection images of the z-stacks. CD31 immunolabeling signal was processed using the despeckle tool in ImageJ to reduce noise, then thresholded by intensity. The %Area covered by thresholded signal in each layer was measured.

For proliferating vascular endothelial cell counts, Hoechst+ nuclei in CD31 immunolabeled vessel structures were identified in the DG layers then categorized as Ki67+ or Ki67-. For density measures, Ki67+CD31+ counts were divided by the area sampled as measured by Zen software. For percent Ki67+ CD31+ cells, the number of double positive cells was divided by the total number of CD31+ cells in that layer to yield a percent measure.

For analysis of NSC distance to vessels, 90 cells/mouse that had a SOX2+ nucleus in the SGZ and a GFAP+ apical process extending through the GCL were identified and the shortest distance from the center of their cell body to the nearest CD31+ vessel structure was measured using the Zen software line measure tool. The identity of the NSCs was confirmed and whether their nucleus was MCM2+ or MCM2-was determined in the full Z-stack images.

For analysis of proliferative cell distance to vessels, 90 cells/mouse that were Ki67+ in the SGZ were identified and their distance to the nearest CD31+ vessel structure was measured using the Zen software line measure tool.

For the analysis of both NSC and Ki67+ location data, the SGZ midline was identified as the border between the GCL and HL and outlined using the Zen software contour tool. The shortest distance to SGZ midline from the center of the cell body of the 90 cells/mouse previously identified was measured using the Zen software line measure tool. Positive distance measurement values were assigned to cells that were closer to the GCL from the SGZ midline. Negative distance measurement values were assigned to cells that were closer to the HL from the SGZ midline. For analysis of Ki67+ location relative to the SGZ midline, GCL-HL border was identified using Hoechst labeling as described above. For the analysis of NSC location relative to the SGZ midline, the SGZ midline was identified as falling directly below the cell dense negative GCL layer in the CD31 labeling.

### Statistical analysis

All statistical tests were conducted using GraphPad Prism, except for 3-way repeated measures ANOVA that was conducted using IBM SPSS Statistics. Data on CD31+ vascular coverage, and density and percent of CD31+ cells that were Ki67+, were analyzed by 2-way repeated measures ANOVA (age x layer) followed by Dunnett’s multiple comparisons test within layer compared to the control age of 9 weeks. Data on distance of MCM2+ NSCs, MCM2-NSCs and Ki67+ proliferative cells to nearest CD31+ vessel were analyzed by ordinary one-way ANOVAs for each cell type followed by Dunnett’s multiple comparisons test for each time point compared to the control age of 9 weeks. Data on location of MCM2+ NSCs, MCM2-NSCs and Ki67+ proliferating cells relative to SGZ midline were analyzed by ordinary one-way ANOVAs for each cell type followed by Dunnett’s multiple comparisons test for each time point compared to the control age of 9 weeks. The overall cell populations of MCM2+ NSCs, MCM2-NSCs and Ki67+ proliferating cells location relative to the SGZ midline were also analyzed by a Kolmogorov-Smirnov test for each time point compared to the control age of 9 weeks. 3-way repeated measures ANOVAs (layer x age x sex) were performed for CD31+ vascular coverage data, and density and percent of CD31+ cells that were Ki67+. 2-way ANOVAs (age x sex) were performed for MCM2+ NSC, MCM2-NSC, and Ki67+ proliferating cells data to determine whether sex effects were present. All data were approximately normally distributed, all statistical tests were two-tailed with α = 0.05. No strong patterns of sex differences were observed in any measure so main data analysis shows data collapsed by sex (Supplemental Fig. S1, S2).

## Data availability

The datasets generated during and/or analyzed during the current study are available from the corresponding author on reasonable request.

## Acknowledgements

This work was funded by R01NS124775 from the National Institute of Neurological Disorders and Stroke to EDK and a National Science Foundation graduate research fellowship program award to ND. We thank Dalia Einstein for assistance with mouse perfusions, and Mia Marcellana and Gwen Sebring for assistance with data collection. Diagrams were created with BioRender.com.

## Author contributions

EDK conceived and designed the study. EDK, ND, AIS, and IC were involved in animal care, experimental procedures, and data collection. EDK, ND, and IC collected data and performed the data analysis. ND and IC wrote the first draft of the manuscript and EDK edited and critically reviewed the manuscript. All authors reviewed the results and approved the final version of the manuscript. ND and IC contributed equally and are co-first authors.

## Additional information

The authors declare no competing interests.

