## Supplementary Fig 3 for "Postnatal development of the dentate gyrus vascular niche"

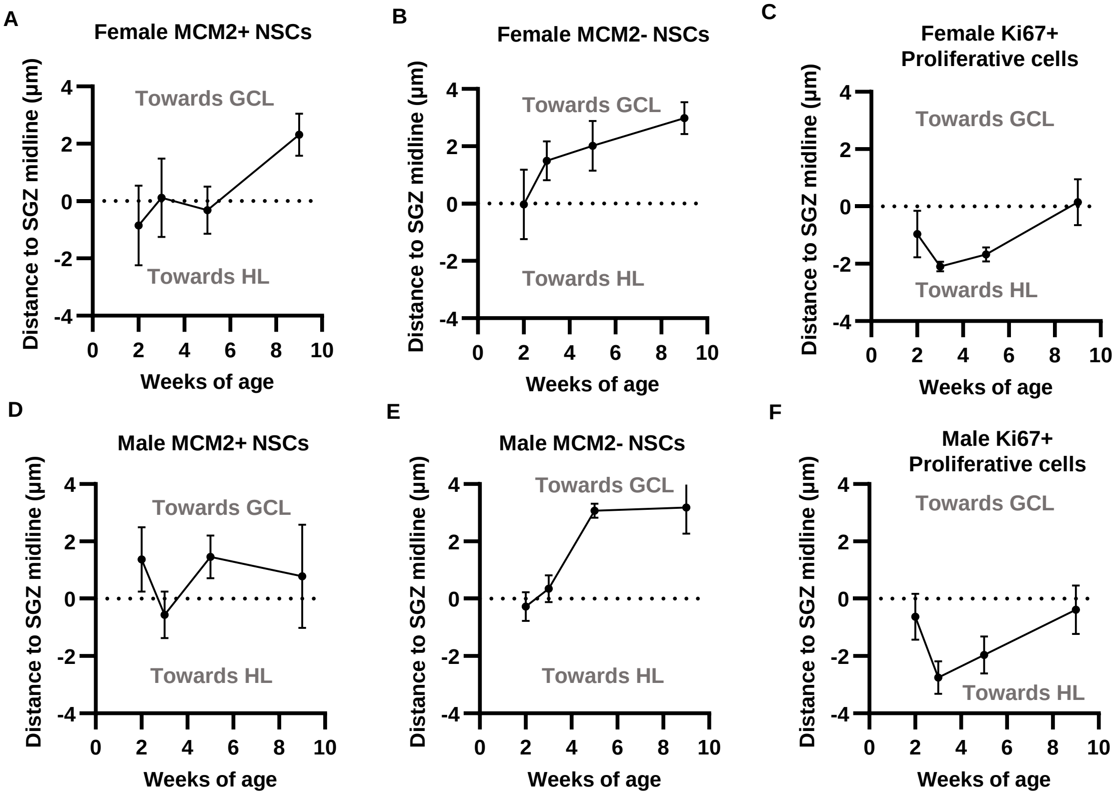


**Fig S3.** Average distance of MCM2- NSCs, MCM2+ NSCs, and Ki67+ Proliferative cells to SGZ midline by sex (**A**), (**D**) MCM2+ NSCs average distance to nearest SGZ midline. 2-way repeated measures ANOVA (age x sex), Age x sex $F_{(3,34)}$ = 1.343, p=0.2768; age $F_{(3,34)}$ = 0.8839, p=0.4592; sex $F_{(1,34)}$ = 0.2393, p=0.6278. (**B**), (**E)** MCM2- NSCs average distance to nearest SGZ midline. 2-way repeated measures ANOVA (age x sex), Age x sex $F_{(3,34)}$ = 0.4429, p=0.7239; age $F_{(3,34)}$ = 7.730, p=0.0005; sex $F_{(1,34)}$ = 0.02913, p=0.8655. **(C),** (**F**) Ki67+ Proliferative cells average distance to nearest SGZ midline. 2-way repeated measures ANOVA (age x sex), Age x sex $F_{(3,34)}$ = 0.1497, p=0.9292; age $F_{(3,34)}$ = 3.438, p=0.0275; sex $F_{(1,34)}$ = 0.2217, p=0.6408**. (A-F)** (n= mice) 2 (n=10, 5M/5F), 3 (n=12, 8M/4F), 5 (n=10, 4M/6F) and 9 (n=10, 5M/5F) weeks.
