## Supplementary Fig 1 for "Postnatal development of the dentate gyrus vascular niche"

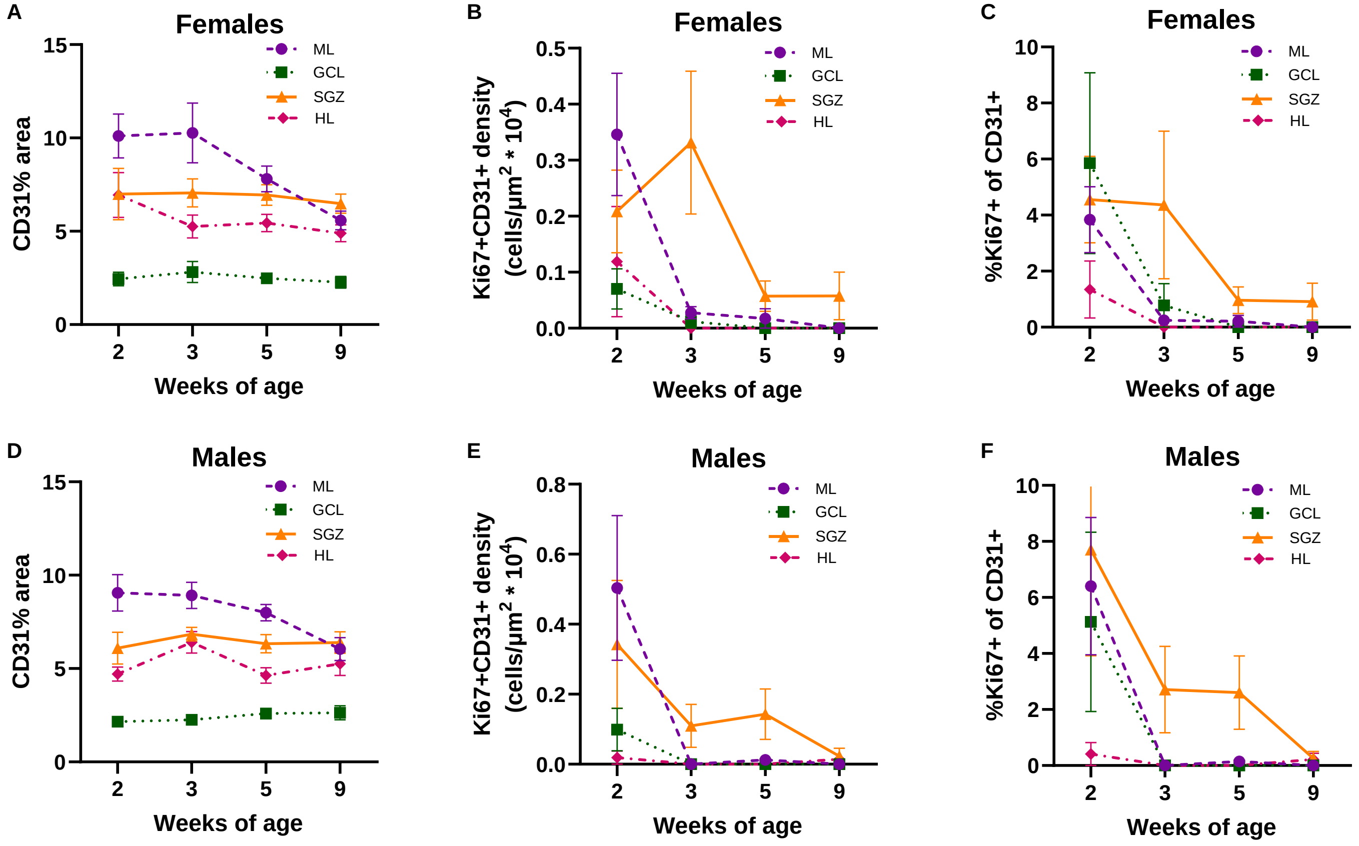


**Fig S1:** Vascular coverage and angiogenesis levels in the postnatal DG by sex. **(A), (D)** Proportion of DG area covered by CD31 immunolabeling. 3-way repeated measures ANOVA age x layer x sex ($F_{9,108}$)=2.56, p=0.02; age x layer ($F_{9,108}$)=9.95, p<0.001; layer x sex ($F_{3,108}$)=0.39, p=0.76; age x sex ($F_{3,36}$)=0.59, p=0.69; age ($F_{3,36}$)=2.08, p=12; sex ($F_{1,36}$)=0.71, p=0.40; layer ($F_{3,108}$)=332.85, p<0.001. **(B), (E)** Density of CD31+/Ki67+ double labeled cells. 3-way repeated measures ANOVA age x layer x sex ($F_{6.18,74.16}$)=1.47, p=0.2; age x layer ($F_{6.18,74.16}$)=5.46, p<0.001; layer x sex ($F_{2.06,74.16}$)=0.36, p=0.70; age x sex ($F_{3,36}$)=0.82, p=0.49; age ($F_{3,36}$)=10.59, p<0.001; sex ($F_{1,36}$)=0.002, p=0.97; layer ($F_{2.06,74.16}$)=13.68, p<0.001. **(C), (F)** Proportion of CD31+ cells co-labeled with Ki67. 3-way repeated measures ANOVA age x layer x sex ($F_{5.41,64.92}$)=0.64, p=0.68; age x layer ($F_{5.41,64.92}$)=2.20, p=0.06; layer x sex ($F_{1.80,64.92}$)=0.41, p=0.65; age x sex ($F_{3,36}$)=0.40, p=0.75; age ($F_{3,36}$)=11.55, p<0.001; sex ($F_{1,36}$)=0.07, p=0.79; layer ($F_{1.80,64.92}$)=8.25, p<0.001. **(A-F)** (n= mice) 2 (n=10, 5M/5F), 3 (n=12, 8M/4F), 5 (n=11, 5M/6F) and 9 (n=11, 5M/6F) weeks
