## Supplementary Fig 2 for "Postnatal development of the dentate gyrus vascular niche"

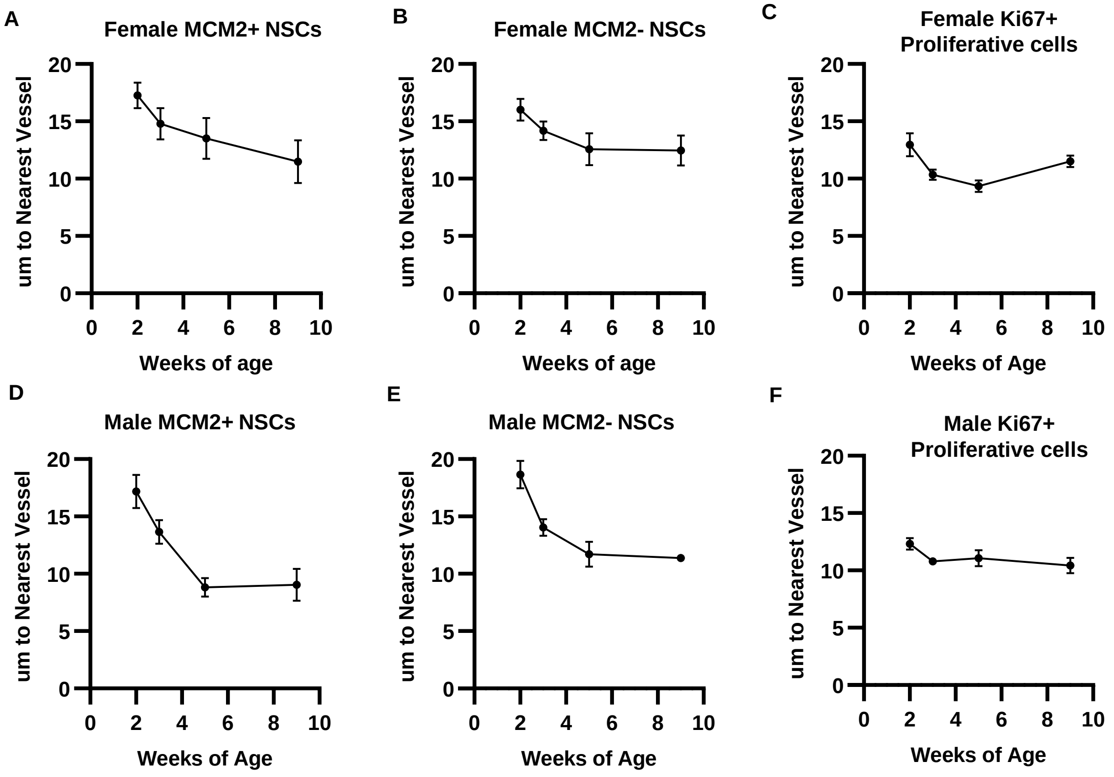


**Fig S2.** Average distance of MCM2- NSCs, MCM2+ NSCs, and Ki67+ Proliferative cells to nearest CD31+ vessel by sex. (**A**), (**D**) MCM2+ NSCs average distance to nearest CD31+ vessel. 2-way repeated measures ANOVA (age x sex), Age x sex $F_{(3,34)}$ = 0.9905, p=0.4089; age $F_{(3,34)}$ = 8.974, p=0.0002; sex $F_{(1,34)}$ = 3.516, p=0.0694. (**B**), (**E**) MCM2- NSCs average distance to nearest CD31+ vessel. 2-way repeated measures ANOVA (age x sex), Age x sex $F_{(3,34)}$ = 1.278, p=0.2975; age $F_{(3,34)}$ = 10.76, p<0.0001; sex $F_{(1,34)}$ = 0.005830, p=0.9396. (**C**), (**F**) Ki67+ Proliferative cells average distance to nearest CD31+ vessel. 2-way repeated measures ANOVA (age x sex), Age x sex $F_{(3,34)}$ = 2.149, p=0.1122; age $F_{(3,34)}$ = 6.412, p=0.0015; sex $F_{(1,34)}$ = 0.07872, p=0.7807. **(A-F)** (n= mice) 2 (n=10, 5M/5F), 3 (n=12, 8M/4F), 5 (n=10, 4M/6F) and 9 (n=10, 5M/5F) weeks.
